# Bio-JOIE: Joint Representation Learning of Biological Knowledge Bases

**DOI:** 10.1101/2020.06.15.153692

**Authors:** Junheng Hao, Chelsea J.-T Ju, Muhao Chen, Yizhou Sun, Carlo Zaniolo, Wei Wang

**Affiliations:** Department of Computer Science, University of California, Los Angeles, 90095, USA; Department of Computer and Information Science, University of Pennsylvania, Philadelphia, 19104, USA

**Keywords:** Biological knowledge bases, representation learning, SARS-CoV-2

## Abstract

The widespread of Coronavirus has led to a worldwide pandemic with a high mortality rate. Currently, the knowledge accumulated from different studies about this virus is very limited. Leveraging a wide-range of biological knowledge, such as gene ontology and protein-protein interaction (PPI) networks from other closely related species presents a vital approach to infer the molecular impact of a new species. In this paper, we propose the transferred multi-relational embedding model `Bio-JOIE` to capture the knowledge of gene ontology and PPI networks, which demonstrates superb capability in modeling the SARS-CoV-2-human protein interactions. `Bio-JOIE` jointly trains two model components. The *knowledge model* encodes the relational facts from the protein and GO domains into separated embedding spaces, using a hierarchy-aware encoding technique employed for the GO terms. On top of that, the *transfer model* learns a non-linear transformation to transfer the knowledge of PPIs and gene ontology annotations across their embedding spaces. By leveraging only structured knowledge, `Bio-JOIE` significantly outperforms existing state-of-the-art methods in PPI type prediction on multiple species. Furthermore, we also demonstrate the potential of leveraging the learned representations on clustering proteins with enzymatic function into enzyme commission families. Finally, we show that `Bio-JOIE` can accurately identify PPIs between the SARS-CoV-2 proteins and human proteins, providing valuable insights for advancing research on this new disease.

## 1 Introduction

The outbreak of COVID-19 (Coronavirus Disease-2019) has infected over 8,000,000 people and caused death for over 430,000 since the end of 2019^1^. Tremendous efforts have been made to discover the infection mechanism of the causative agent, named SARS-CoV-2. One important and urgent task is to understand the mechanism in which viral proteins interact with human proteins. The new findings will enrich the annotation of viral genomes [12] in biomedical knowledge bases (KBs). Constructing and populating such biomedical KBs can significantly improve our understanding of the processes by which SARS-CoV-2 affects different cells in human body and will serve as the foundation for many important downstream applications such as vaccine development [17], drug repurposing [12, 36] and drug side effect detection [37].

In general, biological KBs, often stored as knowledge graphs (KGs), consist of various biological entities, their properties and relations. These KBs can be categorized in different domains, such as gene annotation, functional proteomic analysis, and transcriptomic profiling. Specifically, gene ontology (GO) [10, 16] is the most widely used resource for gene function annotation; STRING [29], PDB [2] and neXtProt [19] collect the knowledge accumulated from functional proteomic analysis; Expression Atlas [25] is a database facilitating the retrieval and analysis of gene expression studies. While those KBs provide the essential sources of knowledge for *in silico* research in the corresponding domains, such domain-specific knowledge is often sparse and costly to apprehend [21, 30]. For example, PPI networks can be far from complete given the information supported by experimental results or suggested by computational inference [14, 21]. Makrodimitris et al. [21] indicate that the numbers of protein-protein interactions in BIOGRID [24] for non-model organisms are far less than expected, specifically, there are only 107 interactions for tomato (*Solanum lycopersicum*) and 80 interactions for pig (*Sus scrofa*). Evidently, relying on the KG from a single domain presents the risk of learning from limited and scarce information.

The stored knowledge is often interrelated across different perspectives. Hence, the missing knowledge in certain KBs can be transferred from other KBs, and thus provide a more comprehensive representation of the biological entities. Taking the protein-protein interaction (PPI) examples of the new SARS-CoV-2 proteins as illustrated in Figure 1, SARS-CoV-2 M protein interacts with a list of 1human proteins, and five of them are involved in the endoplasmic reticulum (ER) morphology process as suggested by the gene ontology annotation (GO:0005783). Similarly, the SARS-CoV-2 ORF3a also interacts with a list of human proteins. Among these proteins, VSP39 and VSP11 are the core subunits of HOPS complex, presenting a binding action as suggested by the STRING database. While aligning the gene ontology annotations of the SARS-CoV-2 M protein as demonstrated in Figure 2, the SARS-CoV M protein presents a similar set of gene ontology annotations, such as “host immune mitigation” and “virion membrane”, suggesting that the side knowledge of gene ontology annotations can facilitate the inference of interactions for related proteins. More generally, the sparse domain information can always benefit from the supplementary knowledge from other relevant domains, therefore calling upon a plausible method to support the fusion and transfer of knowledge across multiple biological domains.

**Fig 1.**
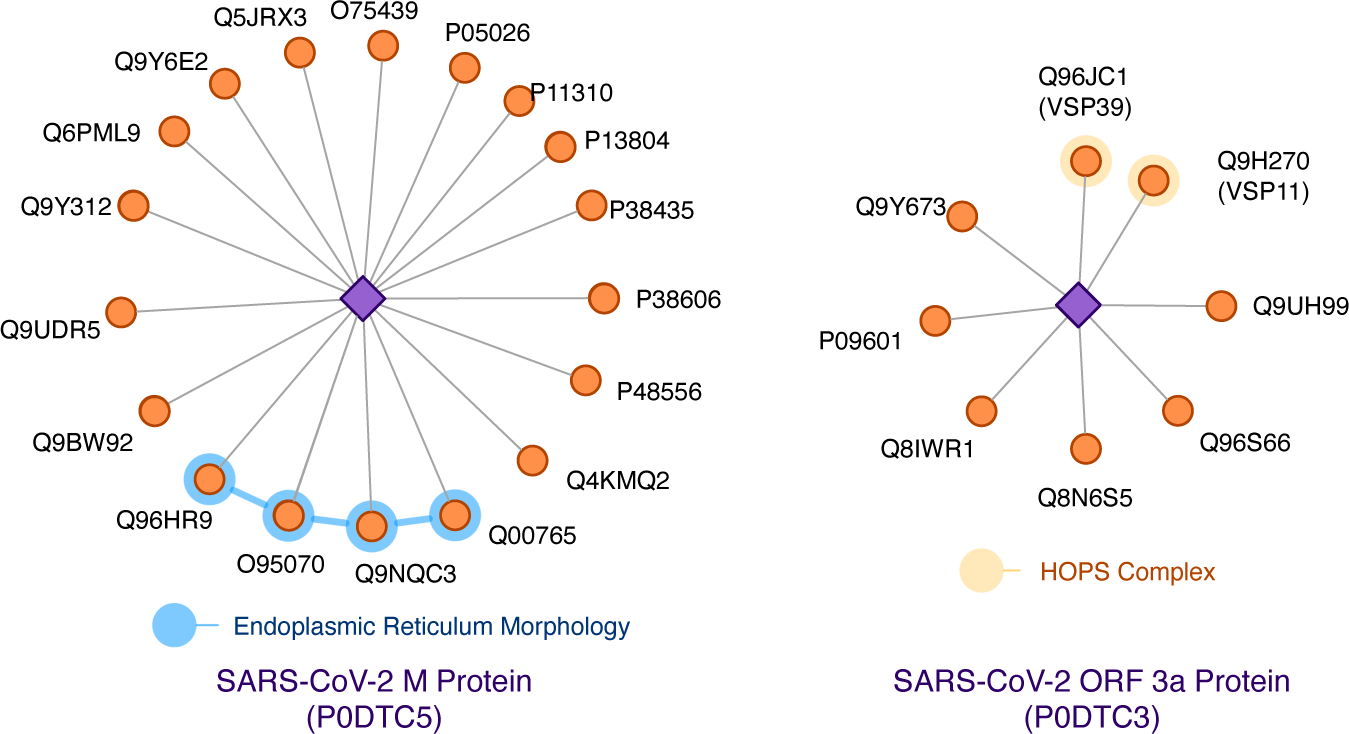
Two examples of SARS-CoV-2-human protein interactions: M protein (left) and ORF3a protein (right). The purple diamonds refers to the viral proteins and the orange circles refer to the high-confidence human protein target. Proteins highlighted in blue are involved in certain biological processes, and proteins highlighted in yellow are arranged in a protein complex.

**Fig 2.**
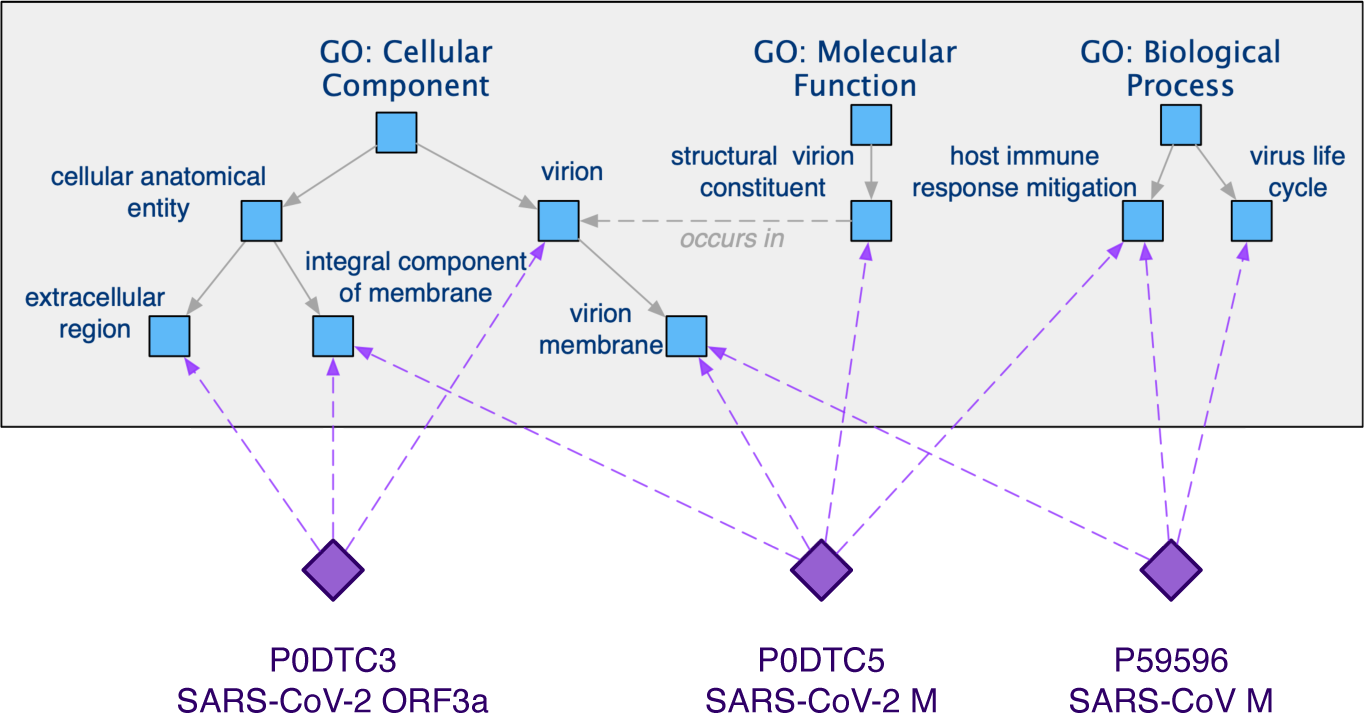
Examples of gene ontology annotation enrichment on three representative SARS-CoV or SARS-CoV-2 proteins, which possess multiple properties across three biological aspects: biological processes, cellular components and molecular functions.

Regardless of the importance and advantages of knowledge fusion across different domains [3, 5], fewer efforts have been devoted to incorporating knowledge from different domains for a specific task in computational biology studies. Onto2vec [27] presents one state-of-the-art learning approach that successfully bridges gene ontology annotations with the protein representation. However, the known PPI information is neglected and not encoded in the obtained protein embeddings.

To combine multiple domain-specific biological knowledge, and facilitate knowledge transfer across different domains, we purpose `Bio-JOIE`, a <u>J</u>oInt <u>E</u>mbedding learning framework for multiple domains of <u>Bio</u>logical KBs. In `Bio-JOIE`, two model components are jointly learned, i.e., a knowledge model characterizes different domain-specific KGs in separate low-dimensional embedding spaces, and a transfer model captures the cross-domain knowledge association. More specifically, the knowledge model encodes the relational facts of entities in each view into the corresponding embedding space separately, with a hierarchy-aware technique designated for the hierarchically-layered domains. Besides, the transfer model seeks to transfer the knowledge between pairs of domains by employing a weighted non-linear transformation across their embedding spaces. In evaluation, we apply the `Bio-JOIE` on several PPI networks with Gene Ontology annotations and the entire gene ontology and evaluate by PPI predictions. We compare `Bio-JOIE` with that of the state-of-the-art representation learning approaches on multiple species, including SARS-CoV-2-Human PPIs, with different model settings. Our best `Bio-JOIE` outperforms alternative approaches by 7.4% in PPI prediction.

Our contributions are 4-fold. First, we construct a general framework for learning representations across different domain-specific KBs, including the dynamically changing SARS-CoV-2 KB. Second, we emphasize and demonstrate that cross-domain representation learning by the proposed `Bio-JOIE` can improve the inference in one domain by leveraging the complementary knowledge from another domain. Extensive experiments on different species confirm the effectiveness of cross-domain representation learning. Third, `Bio-JOIE` also demonstrates cross-species transferability to improve PPI predictions among multiple species by knowledge population from gene ontology. Fourth, the protein representations learned from `Bio-JOIE` can be leveraged for different tasks. Specifically, we show that the protein embeddings trained on PPI network and gene ontology present the potential to better group enzymes into different enzyme commission families.

## 2 Materials and Method

In this section, we present the proposed method to support representation learning and cross-domain knowledge transfer on biological KBs. Without loss of generality and aligned with the evaluation of the proposed `Bio-JOIE`, we refer two domain-specific KGs in the following section to PPI networks and the gene ontology graph. We begin with the formalized descriptions of the materials and tasks.

### 2.1 Preliminary

#### Materials

A typical biological KB can be viewed as relational data that are presented as an edge-labeled directed graph *G*, which is formed with a set of entities (e.g. proteins) *ɛ* and a set of relations (e.g. interaction types) *R*. A triple (*s, r, t*) ∈ *G* represents a *r* ∈ *R* typed relation between the source and target entities *s, t* ∈ *ɛ*, As stated, we continue with the modeling on KGs of two domains, PPI and gene ontology. For example, in the PPI network, a triple (`FBgn0011606`, `binding`, `FBgn0260855`) simply states the fact that two proteins (from fly) have binding interaction; and in gene ontology, a triple (`GO:0008152`, `is a`, `GO:0008150`) similarly represents that `GO:0008152` (a unique identifier of “metabolic process”) is one subclass of `GO:0008150` (a unique identifier of “biological process”). Our model seeks to capture the protein information in the triples (*s_p_*, *r_p_*, *t_p_*) of PPI graph *G_p_* in a *k_p_*-dimensional embedding space, where we use boldfaced notations such as **s**_*p*_, **r**_*p*_, **t**_*p*_ ∈ ℝ^*kp*^ to denote the embedding representation. Similarly, gene ontology is another graph *G_o_* formed with a set of GO terms *ɛ_o_* and a set of semantic relations *R_o_*. The triple (*s_o_*, *r_o_*, *t_o_*) ∈ *G_o_* identifies a semantic relation of GO terms, while we also observe hierarchical substructures formed by “subclass” or “is_a” relation as the aforementioned example. The gene ontology is embedded in another space ℝ^*ko*^, such that *k_p_* and *k_o_* may not be equivalent. We use (*o, p*) ∈ *A* to denote a *GO term annotation* where a GO term *o* ∈ *E_o_* describes a protein *p* ∈ *E_p_* of its corresponding functionality, and *A* denotes the set of such associations. As introduced in Section 1, we consider SARS-CoV-2-Human interaction as a similar (but significantly smaller) KBs with the same structures as *G_p_*, which serves as an extension of human PPI networks.

#### Tasks

To validate the learned embedding of biological entities (proteins and GO terms in this context), we address the following two tasks. (i) *PPI type prediction* aims at predicting the interaction type between two interacting proteins, including *SARS-CoV-2 related PPIs*; (ii) *Protein clustering and family identification* aims at clustering the existing proteins and helps identify the clusters based on Enzyme Commission (EC) numbers.

#### Methods

The model architecture of `Bio-JOIE` is shown in Figure 3. The proposed `Bio-JOIE` jointly learns two types of model components to connect the two views of structured knowledge. Knowledge models are responsible for representing the relational knowledge of PPI and that of GO term into two separate embedding spaces ℝ^*kp*^ and ℝ^*ko*^ a by using KG embedding and hierarchy-aware regularization. On top of that, a transfer model learns a transformation to connect between the representations of GO term relation facts and PPI based on partially provided GO term assignments. In particular, we investigate weighted *transfer techniques* to better capture the knowledge transfer, for which the weights reflect the specificity of the assigned GO term to a protein. The following of this section describes the model components and the learning objective of `Bio-JOIE` in detail.

**Fig 3.**
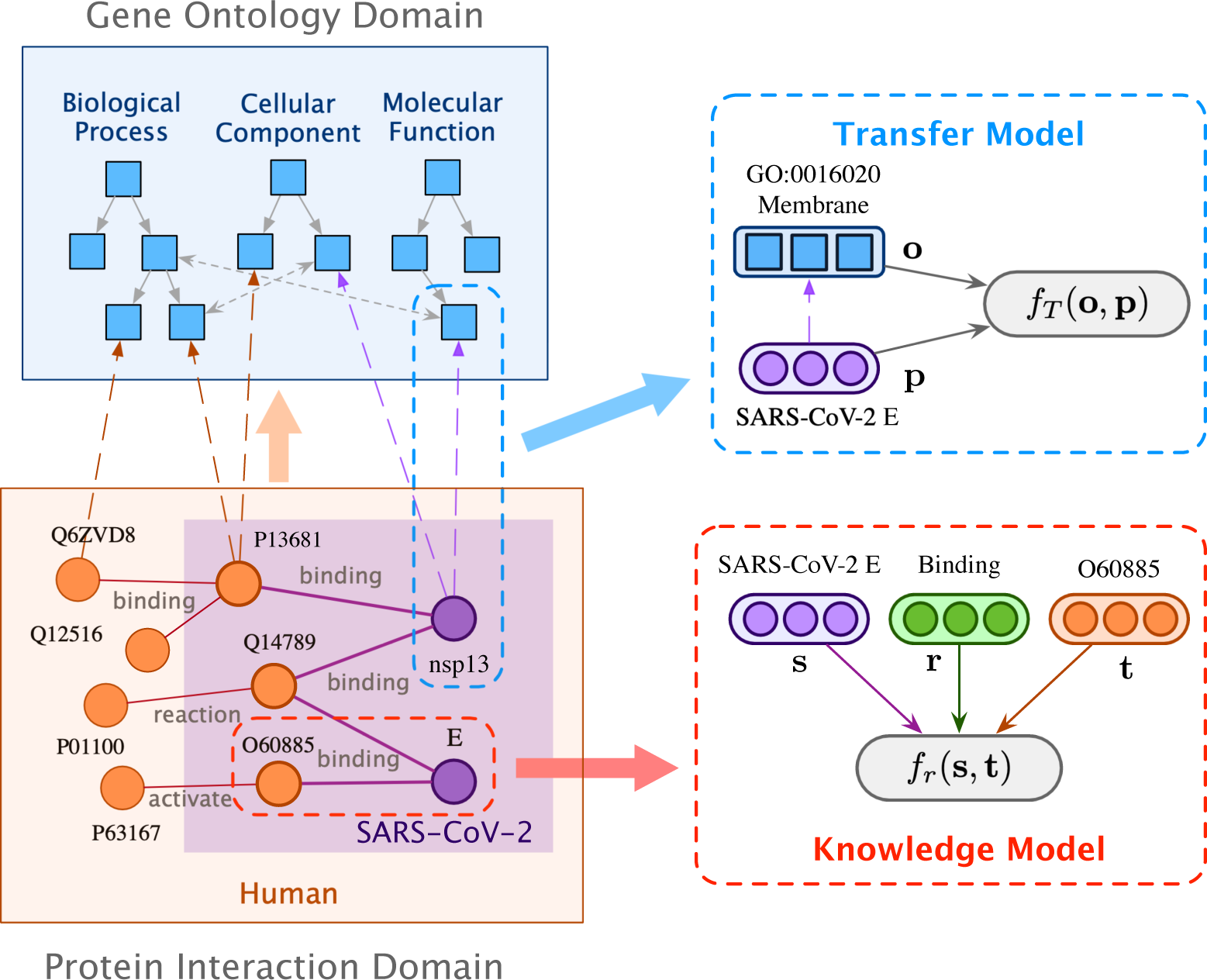
Model architecture of `Bio-JOIE`. The Knowledge Model seeks to encode relational facts in each domain respectively (such as proteins and gene ontology). Meanwhile, the Transfer Model learns to connect both domains and enable knowledge transfer across protein and gene ontology.

**Fig 4.**
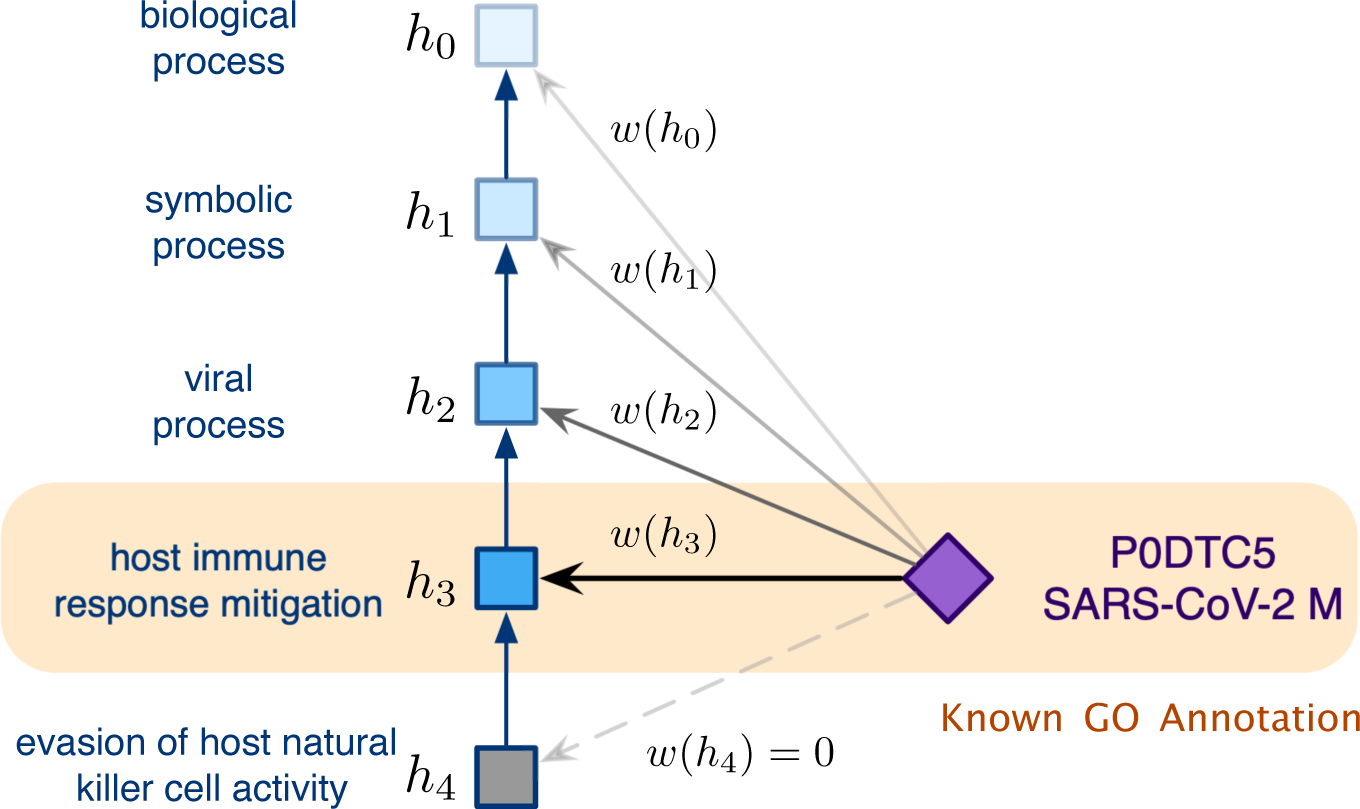
Explanation of weighted transfer model for modeling hierarchical gene ontology.

### 2.2 Knowledge Model

The knowledge models seek to characterize the semantic relations of GO terms and PPI information into separate embedding spaces. In each embedding space, the inference of relations or interactions is modeled as specific algebraic vector operations. As mentioned, the two views of gene ontology and PPI are embedded to separate embedding spaces.

To capture a triple (*s, r, t*) from either of the two domains, a cost function *f*_*r*_ (*s, t*) is provided to measure its plausibility. A lower score indicates a more plausible triple. We can adopt multiple vector operations in the defined embedding space with three representative examples defined as follows, i.e. translations (TransE [4]), Hadamard product [33] and circular correlation (HolE [23]). The cost functions are given as follows, where the symbol ◦ denotes Hadamard product, and ∗ : ℝ^*d*^ × ℝ^*d*^ → ℝ^*d*^ denotes circular correlation defined as 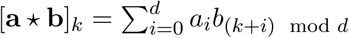.

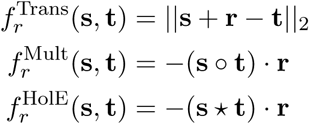

Since most of the relations in PPI networks are symmetric (such as binding and catalysis), we apply the Hadamard product based function. The learning objective of a knowledge model on a graph *G* is to minimize the following margin ranking loss,

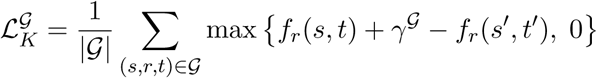

where *γ^G^* is a positive margin, and a negative sample (*s’, r, t’*) ∈ *G* is created by randomly substituting either *s* or *t* using Bernoulli negative sampling [32]. With regard to the two domains of relational knowledge (proteins and gene ontology) *G*_*p*_ and *G*_*o*_, we denote the learning objective losses 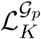 and 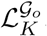.

#### Hierarchy-aware Encoding Regularization

As mentioned in Section 2.1, it is observed that some ontological knowledge can form hierarchies [8], which is typically constituted by a relation with the implicit hierarchical property, such as “subclass of”, as substructures. In gene ontology, more than 50% of the triples have such relations. To better characterize such hierarchies, we model such substructures differently from the aforementioned DistMult and many others by adding hierarchy regularization. More specifically, given entity pairs (*e*_*l*_, *e*_*h*_) ∈ *S* where *e*_*l*_ is a subclass of *e*_*h*_, we model such hierarchies by minimizing the distance between coarser concepts and associated finer concepts in embedding space. Hence, the loss is simply defined as

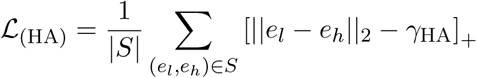

where [*x*]_+_ = max{*x*, 0} and *γ*HA is also a positive margin parameter. This penalizes the case where the embedding of *ɛ*_*l*_ falls out the *γ*HA-radius neighborhood centered at the embedding of *e_h_*.

*Relation Inference* Given the learned embeddings and a pair of query proteins ((*p*_1_, *p*_2_)), we can predict the most plausible interaction type *r* by selecting the optimal *f*_*r*_ (*p*_1_, *p*_2_) score. We can also provide predictions for possible protein targets given the query of the subject protein and specific interaction type (*p, r*, ?*t*) by populating the selection proteins with top score *f*_*r*_ (*p, t*) from the knowledge model. Details about each task are curated in Section 3.3 and 3.5.

### 2.3 Transfer Model

The transfer model learns to connect between the above two relational embedding spaces via a non-linear transformation. The transformation is induced based on the GO term assignments, towards the goal to collocate the associated GO terms and proteins in an embedding space after transformation. Hence, the affinity of embedding structures of gene ontology and PPIs can be captured. This allows the relational knowledge to transfer across and complement the learning and inference on both domains.

Given each GO term assignment (*o, p*) ∈ *A*, following function *f_T_* (*o, p*) measures the plausibility of the transformation that is favored to be minimized.

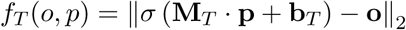

**M**_*T*_ ∈ ℝ^*ko×kp*^ thereof is a weight matrix and **b**_*T*_ ∈ ℝ^*kp*^ is a bias vector. *σ* is either the identify function, or a non-linear function as tanh, the latter thereof aim at smoothing the transformation with additional non-linearity.

#### Basic Transfer Model

The basic strategy to learn the transfer model is to treat each GO term assignment evenly, and thereby minimizing the following learning objective loss.

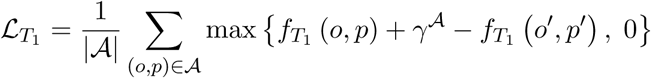

(*p’, o’*) *A* thereof is a negative sample by randomly substituting *p’*, and *γ^A^* is a positive margin.

#### Weighed Transfer Model

Since some ontological knowledge, such as gene ontology, may form hierarchical structures, where GO terms in lower levels typically describe more specified gene functionality. During the characterization of associations between GO terms and proteins, in contract to general GO terms, more specified GO terms necessarily carry more precise descriptions to the proteins. Hence, an improved transfer model weights among GO term associations to a protein for the purpose of more attentively capturing those with more specific GO terms. Let *ω*(*o*) be a weight is specifically assigned to *o*, the objective of the weighted transfer model is to minimize the following loss,

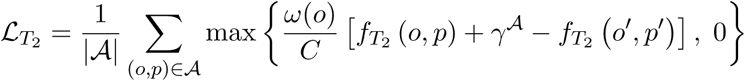

where *C* is a normalizing constant to constrain that 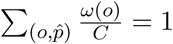 for a specific protein 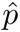.

Exemplarily, there could be several ways to calculate the association weight.

#### Level-based weight

The level of the node in one hierarchical taxonomy is a natural indicator of its specificity. Accordingly, the weight can be defined as,

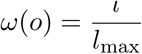

where *l* is the term’s current depth and *l*_max_ is the maximum length of the associated branch in the gene ontology DAG.

#### Degree centrality weight

A small node’s degree centrality in the graph roughly reflects its specialty and we apply

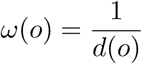

as the balance factor for different GO term specialty.

In practice, incorporating a specificity-based weight to the transfer model essentially enhances the inference in the protein domain, as we have observed in the evaluation in Section 3. However, the above weight options generally yield similar performance gain, and we fix the weight option as the level-based weight in our experimental setting.

### 2.4 Joint Learning Objectives

`Bio-JOIE` jointly learns two knowledge models respectively for GO term relations and PPIs, and a transfer model to support knowledge transfer between these two. Therefore, the joint learning objective minimizes the following loss:

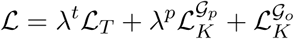

*λ^p^* and *λ^t^* are two positive hyperparameters. We use Adam [18] to optimize the learning objective loss. The learning process uses orthogonal initialization [26] to initialize the weight matrix, and Xavier normal initialization [11] for vector parameters. A normalization constraint is enforced to keep all embedding vectors of GO terms and proteins on unit hyper-spherical surfaces, which is to prevent the non-convex optimization process from collapsing to a trivial solution where all vectors shrink to zero [4, 13, 20, 33].

Note that `Bio-JOIE` is suitable for joint representation learning on proteomic knowledge of different species. In this protein-GO example, the proteins of these species are significantly different from each other. However, they share the same set of annotations in the GO domain. Therefore, More specifically, if we have multiple PPI networks *G_i_*, *i* = 1, 2,. . ., *m* where *m* denotes the number of independent species, *n* knowledge models are trained respectively. Consequently, one unique transfer model is also trained to facilitate the protein-GO knowledge transfer regarding each species. The learning objective on the multi-species setting is changed accordingly as,

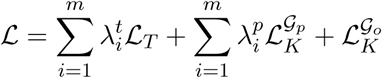

with the assumption that the knowledge model for gene ontology remains unchanged.

In addition to joint learning on multiple species, `Bio-JOIE` can also be re-trained from new observations of PPIs. For example, suppose newly discovered SARS-CoV-2-Human PPI knowledge extends the original human PPI networks, we can fine-tune the `Bio-JOIE` from the saved model and obtained embeddings, by only optimizing the `Bio-JOIE` on the new triples and hence fast obtain representations for all new proteins, without a long time for retraining the `Bio-JOIE` from scratch.

## 3 Results

In this section, we evaluate the embeddings learned from `Bio-JOIE` with two groups of tasks: PPI type prediction (Section 3.3) and protein clustering based on enzymatic functions (Section 3.4). Furthermore, we provide an extensive case study in Section Section 3.5 on SARS-CoV-2 related PPI prediction and classification.

### 3.1 Dataset

The protein-protein interactions for yeast (*Saccharomyces cerevisiae*), fly (*Drosophila melanogaster*), and human (*Homo sapiens*) are collected from the STRING [29] database. There are seven types of interactions annotated in the STRING database. To preserve a balanced and sufficient number of cases in each class, we randomly choose the protein pairs from four types of interaction: activation, binding, catalysis, and reaction. In total, there are 21704, 10000, 36400 pairs of proteins for yeast, fly, and human, respectively; each type contains roughly the same number of interactions. Table 1 summarizes the PPI information for each species. Note that, the human PPI dataset does not contain the virus-generated proteins, but the set partially overlaps with the virus-human pan-PPI networks.

**Table 1.** Statistics of PPI networks and associated GO annotations.

| Species | # Proteins | # PPI Triples | # GO Annotations |
| --- | --- | --- | --- |
| Yeast | 3,736 | 21,704 | 191,801 |
| Fly | 3,826 | 10,000 | 87,807 |
| Human | 8,204 | 36,400 | 102,759 |

The gene ontology annotations for each protein are extracted from gene ontology Consortium [10], including all three biological aspects: biological process (BP), cellular components (CC), and molecular function (MF). Table 2 summarizes the number of relations between proteins and GO terms. The relations between GO terms include *is-a*, *part-of*, *has-part*, *regulates*, *positively-regulates*, and *negatively-regulates*.

**Table 2.** Statistics of three aspects in the gene ontology: biological processes (BP), cellular components (CC) and molecular functions (MF).

| Aspects | BP | CC | MF |
| --- | --- | --- | --- |
| # GO entities | 5744 | 1,147 | 1,764 |
| # GO triples | 19,021 | 2,116 | 2,190 |
| # Protein-GO annotations (yeast) | 72,956 | 58,729 | 60,116 |
| # Protein-GO annotations (fly) | 44,605 | 24,550 | 18,652 |
| # Protein-GO annotations (human) | 42,899 | 32,929 | 26,931 |

For the SARS-CoV-2 dataset, we collect the latest virus-protein interaction from BioGrid^2^ and the limited GO annotations for SARS-CoV-2 from Gene Ontology Consortium^3^, as last updated on early April. In summary, there are 26 SARS-CoV-2 generated proteins and 332 human proteins presenting the evidence of viral-human protein interactions as suggested by Gordon et al. [12]. The selection is based on a high MIST score and a low SAINTexpress BFDR from Affinity Capture-MS. Out of the same experiment, we select 1131 viral-human protein pairs with MIST scores lower than 0.01 as our negative samples. The 26 SARS-CoV-2 generated proteins are annotated with 282 GO terms. In addition to SARS-CoV-2, BioGrid also includes 30 viral proteins from SARS-CoV and MERS-CoV, which are two similar contagious viruses causing respiratory infection. These 30 viral proteins are annotated with 630 GO terms, and display 326 interactions with human proteins. All processed datasets are available at https://www.haojunheng.com/project/goterm.

### 3.2 Baselines

We compare `Bio-JOIE` with the most applicable state-of-the-art approach, Onto2Vec [27], on learning the representation of proteins. Onto2Vec considered the annotation from gene ontology for representation learning. In addition, we compare `Bio-JOIE` with a simpler setting, `Bio-JOIE`-NonGO, where we only consider the single-domain knowledge of PPI.

#### Onto2Vec, Onto2Vec-Parent, Onto2Vec-Ancestor

Onto2Vec utilizes the annotation information from gene ontology to create pairwise context and apply Word2Vec [22] to generate protein and GO term embeddings. Its schema allows the model to learn the representation of proteins and GO terms simultaneously. The proposed setting of Onto2Vec only includes the direct relationship between a protein and a GO term. In this experiment, we explicitly include the relationship between a protein and the parents of the annotated GO terms, named *Onto2Vec-Parent*, and the ancestors of the annotated GO terms, named *Onto2Vec-Ancestor*.

#### Onto2Vec-Sum, Onto2Vec-Mean

To examine the effect of Onto2Vec on learning the protein representation from a single domain, i.e. gene ontology, we remove the relations between proteins and GO terms during the learning process. The representation of a protein is then computed by either summing up the embeddings of all the associated GO terms (*Onto2Vec-Sum*), or taking the average of the embeddings of those GO terms (*Onto2Vec-Mean*).

#### OPA2Vec

Based on Onto2Vec, OPA2Vec further learns the protein and GO term embeddings by leveraging meta-data (labels, synonyms, etc), which better characterize GO terms. `Bio-JOIE` **(NonGO)**. As opposed to considering the knowledge from a single domain of gene ontology, we adopt `Bio-JOIE` to consider only the knowledge from Protein-Protein Interaction. In this approach, all the gene ontology annotations and the gene ontology graph are neglected, and thus is reduced to a knowledge model. We only use the knowledge model in Section 2.2, where the protein embeddings are solely learned from PPI networks by the original KG embedding technique, DistMult. We refer to this approach as “Non-GO”.

It is worth mentioning that the goal of Onto2Vec and OPA2Vec is to learn the protein representation; therefore, to adapt for the task of PPI prediction, we concatenate the embeddings of each pair of proteins and train a multi-class classifier to predict the PPI type for a given pair of query proteins. We examine the performance with four different classifiers: logistic regression (LR), support vector machine (SVM), random forest (RF), and neural networks (MLP). The evaluation is conducted with five-fold cross-validation. Similar settings apply to all Onto2Vec variants and OPA2Vec. On the contrary, our proposed model equips with relational modeling and outputs PPI predictions by selecting the most plausible relation type. As a result, we do not need an additional classifier for `Bio-JOIE` and `Bio-JOIE`-NonGO.

### 3.3 PPI Type Prediction on Multiple Species

We examine how effectively `Bio-JOIE` leverages gene ontology to predict protein-protein interaction types. To do so, we first evaluate the performance on three organisms separately: human, yeast, and fly. Then we study the contribution of the three aspects in gene ontology, i.e. biological process (BP), cellular component (CC), and molecular function (MF), on predicting the type of PPI. Specifically, we provide an analysis on how the knowledge from Gene Ontology contributes to PPIs in different species.

#### Experimental setting

We first separate the PPI triples into approximately 70% for training, 10% for validation and 20% for testing. For hyperparameters with the best performance from the validation set, we select dimension *dp* = *do* = 300 and margin parameters *γG* = 0.25, *γA* = 1.0 and *γ*HA = 1.0. Two weight factors in the joint learning objective are set as *λ_p_* = 1.0, *λ_t_* = 1.0. We use DistMult for the knowledge model in Section 2.2, with hierarchy-aware regularization and the level-weighted transfer model (Section 2.3) deployed. For simplicity, the reported `Bio-JOIE` adopt the same settings if not specifically explained. The number of epochs in training on all settings is limited to 150. For evaluation, we aim at predicting the correct interaction type, given pairs of proteins in the test set. We conduct a 5-fold cross validation for `Bio-JOIE` and all baselines, and report the average and standard deviation of accuracy. The best-performing classifier is RF for OPA2Vec and most of the Onto2Vec variants. The only exception is to apply MLP for Onto2Vec-Ancestor on fly.

#### Results

The results for PPI type prediction are shown in Table 3. We observe that our best `Bio-JOIE` variant outperforms `Bio-JOIE`-NonGO by 7.4% on average for all three species. This observation directly shows that gene ontology KG provides complementary knowledge for proteins. Subsequently, Gene Ontology annotations benefit the learning of protein representations and better predict the interaction types between proteins. Compared to other baselines, it is observed that `Bio-JOIE` notably outperforms Onto2Vec-Ancestor with an average increase of 7.4% on the prediction accuracy, and a relative gain of 9.0% on average of all three species. This observation is due to the advantage that `Bio-JOIE` better leverages the complementary knowledge from PPI to enhance the PPI prediction. As mentioned in Section 3.2, Onto2Vec does not utilize the PPI information into protein embedding learning. Instead, it obtains embeddings based on the aggregated semantic representations of GO terms. It requires additional classifiers for PPI type prediction given pretrained protein embeddings. In contrast, `Bio-JOIE` jointly learns protein representations from both the knowledge model that captures the structured information of known PPIs, and the transfer model that delivers the annotations of GO terms. Also, we observe that `Bio-JOIE`-Weighted achieves better results than `Bio-JOIE`, with a relative performance gain of 2.5%. We hypothesize that such gain is attributed to specificity modeling in the transfer model which distinguishes more specific and informative GO terms from other general GO terms and assigns a higher weight, which selectively learns the alignments between two domains. In terms of different species, we also observe that `Bio-JOIE` achieves a higher PPI prediction accuracy on yeast compared to human and fly. The possible reason is that the yeast interaction network is denser, such that 0.30% of the protein pairs are known to interact, compared to human (0.13%) and fly (0.11%), which indicates that yeast is possibly well studied. OPA2Vec claims to be an improved version of Onto2Vec. Similar to Onto2Vec, it only considers the direct relationship between a protein and a GO term, without parents and ancestors. We find that OPA2Vec performs slightly better than Onto2Vec on Yeast and Fly, but worse on Human. In addition, OPA2Vec falls short when compared to any of the `Bio-JOIE` variants, indicating that incorporating the metadata of GO terms is insufficient for protein representation learning.

**Table 3.** PPI type prediction accuracy (%) evaluated on yeast, fly and human species.

| Model | Yeast | Fly | Human |
| --- | --- | --- | --- |
| Onto2Vec | 76.41 ± 0.73 | 70.85 ± 0.85 | 77.97 ± 0.46 |
| Onto2Vec-Parent | 80.79 ± 0.66 | 75.46 ± 1.11 | 74.90 ± 0.46 |
| Onto2Vec-Ancestor | 86.31 ± 0.42 | 80.31 ± 0.92 | 78.73 ± 0.46 |
| Onto2Vec-Sum | 76.38 ± 0.83 | 72.84 ± 1.13 | 72.53 ± 0.73 |
| Onto2Vec-Mean | 77.95 ± 0.81 | 74.38 ± 1.13 | 73.47 ± 0.80 |
| OPA2Vec | 79.88 ± 0.74 | 74.45 ± 0.97 | 72.04 ± 0.58 |
| Bio-JOIE-NonGO | 83.65 ± 0.92 | 77.58 ± 1.07 | 76.10 ± 0.87 |
| Bio-JOIE | 87.15 ± 1.15 | 84.56 ± 0.81 | 81.42 ± 0.62 |
| Bio-JOIE-Weighted | **90.12 ± 1.21** | **85.55 ± 1.57** | **83.89 ± 0.92** |

It is noteworthy that unlike Onto2Vec, which achieves its best performance with the help of full gene ontology (i.e. Onto2Vec-Ancestor), our `Bio-JOIE` model can utilize only the GO terms that are directly annotated with the proteins to accomplish the highest accuracy score. This also makes `Bio-JOIE` training processes more time efficient. We hypothesize that for `Bio-JOIE` in the PPI type prediction task, GO terms that are directly related to associated proteins with high specificity are sufficient for the transfer model to model the protein-GO association in the embedding spaces. In contrast, Onto2Vec needs entire structured information of GO terms for its word2vec module to construct an exhaustive context of protein features.

We further explore the effects of three different aspects of gene ontology in predicting the types of PPIs. To achieve this, we train `Bio-JOIE` in settings where only specific aspects of gene ontology annotations are used. Results are shown in Table 4, in which BP, CC and MF respectively refer to the cases where GO terms of *biological processes*, *cellular components* and *molecular functions* are used. “BP + CC” denotes that the GO terms from both biological processes and cellular components are included in training. We observe that `Bio-JOIE` performs the best with GO terms from all aspects (full gene ontology). This phenomenon is consistent among all three species, indicating that the protein representations are more robust when learning from a more enhanced knowledge graph. It is also interesting to see that the accuracy of the task is generally higher when we include the GO terms from biological processes. This leads to 2.61% improvement in accuracy over CC, and at least 2.13% of improvement over MF when evaluating individually. In the two-aspect evaluation, “BP+CC” is in average leads to 0.7% better accuracy than “CC+MF”. This is attributed to the fact that BP is the largest group in the gene ontology, containing more entities and relational facts. Consequently, `Bio-JOIE` achieves the best performance with all three aspects of gene ontology annotations incorporated. This indicates that the characterization of PPIs benefits from more comprehensive gene ontology annotations.

**Table 4.** Comparison of PPI prediction accuracy of `Bio-JOIE` on three different aspects of gene ontology.

| # | Aspects | Yeast | Fly | Human |
| --- | --- | --- | --- | --- |
| 1 | BP | 0.8794 | 0.8402 | 0.8153 |
|  | CC | 0.8499 | 0.8272 | 0.8054 |
|  | MF | 0.8539 | 0.8386 | 0.8165 |
| 2 | BP+CC | 0.8717 | 0.8473 | 0.8271 |
|  | BP+MF | 0.8673 | 0.8471 | 0.8163 |
|  | CC+MF | 0.8569 | 0.8466 | 0.8170 |
| 3 | All-GO | **0.9012** | **0.8555** | **0.8389** |

In addition to joint learning from two different domains (i.e. GO terms and PPIs), as mentioned in Section 2.4, `Bio-JOIE` can be trained to capture PPIs for multiple species with several species-specific knowledge models, along with transfer models that bridge the universal gene ontology. To validate the benefit of joint learning on multiple species together, we consider three following configurations of `Bio-JOIE`: (i) the “multi-way” setting uses one unique knowledge model and one transfer model to the universal gene ontology for each species; (ii) the “concat” setting uses one unified knowledge model to capture all species of PPIs, together with one transfer model to learn protein-GO alignments, that is, simply concatenate all PPI triples and all gene ontology annotations of proteins in multiple species; (iii) the “single” setting trains separately on each species, which is exactly the same as in the setting in Table 3. We summarize the results in Table 5. It is observed that the “multi-way” setting can slightly improve PPI performance in comparison to the “ single” setting that trains separately on each species. Also in the “concat” setting with one shared transfer model and knowledge model, the performance significantly drops with a 2.8% decrease of accuracy on average compared to the “single” setting. Such results suggest that each species has unique patterns of PPIs, such differences are better differentiated in separate embedding spaces. Hence, the multi-way setting better encodes the species-specific knowledge and model, which helps the type prediction of PPIs for each species by `Bio-JOIE` that are jointly trained on multiple species.

**Table 5.** PPI type prediction accuracy on different configurations of multi-species joint learning.

| Model | Yeast | Fly | Human |
| --- | --- | --- | --- |
| Bio-JOIE (single) | 0.9012 | 0.8555 | 0.8389 |
| Bio-JOIE (concat) | 0.8795 | 0.8282 | 0.8028 |
| Bio-JOIE (multi-way) | **0.9062** | **0.8638** | **0.8426** |

### 3.4 Identifying Protein Families And Enzyme Commission Based Clustering

Besides inferring PPI types, the embedding representations of proteins can also be used to identify potential protein families based on their functions. This can be achieved by performing clustering algorithms on the learned protein embeddings.

The Enzyme Commission number (EC number) defines a hierarchical classification scheme that provides the enzyme nomenclature based on enzyme-catalyzed reactions. The top-level EC numbers contain seven classes: oxidoreductases, transferases, hydrolases, lyases, isomerases, ligases, and translocases. In this experiment, we select 1340 yeast proteins in total with enzymatic functions. We learn the protein representations using all the triples of PPI networks and the annotation from gene ontology and evaluate the learned representations of these proteins by performing the k-means clustering algorithm to group them into seven non-overlapping clusters. These clusters are compared with the top-level of enzyme commission classification. Purity score is reported as evaluation metrics.

The evaluation of the clustering results is shown in Table 6. `Bio-JOIE` achieves the best clustering performance on yeast by a relative increase of 9.7%, which demonstrates that `Bio-JOIE` has the good model capability to representation learning and empirically show the validity of the learned embeddings to measure the similarity. We hypothesize that `Bio-JOIE` better incorporates protein annotation resource and utilizes the complementary knowledge in the gene ontology domain, while `Bio-JOIE` also captures PPI information and encode it into protein embeddings. This in the end results in comprehensive representations for proteins and helps to identify protein EC classes by clustering.

**Table 6.**
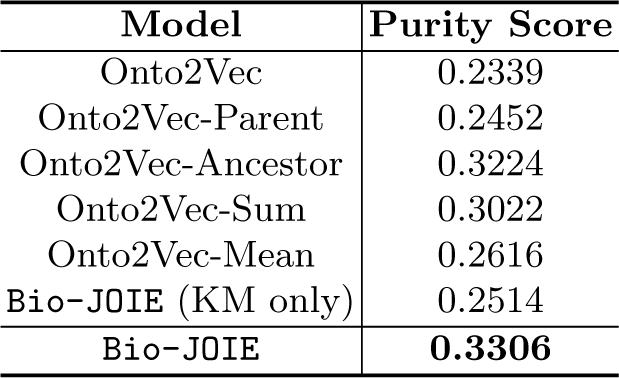
Results of top-level EC clustering by K-means on learning selected yeast protein embeddings.

### 3.5 Case Study: SARS-CoV-2-Human Protein Target Prediction

The COVID-19 pandemic requires many efforts and attentions from scientists of different fields. However, there is very limited knowledge of the molecular details of SARS-CoV-2. In this sub-section, we apply `Bio-JOIE` to gain more insights of the PPI network between SARS-CoV-2 and human proteins. Specifically, we explore the potential of `Bio-JOIE` on predicting whether a pair of human and SARS-CoV-2 proteins interact or not. This is modeled as a binary prediction task. Correspondingly, results from the binary predictions can serve as a guide to identify the targeted proteins by SARS-CoV-2. We first use the known interactions between these two species to validate the effectiveness of `Bio-JOIE`. These interactions are experimentally verified as described in Section 3.3. In this setting, we particularly study the contribution of the knowledge of other closely related viruses (SARS-CoV and MERS) on supporting PPI prediction. We also show the high-confidence candidates of targeted human proteins predicted by `Bio-JOIE` for four selected SARS-CoV-2 proteins.

#### Experimental setting

In this experiment, we randomly split the known positive human-virus PPIs into train and test sets with a ratio of 80% and 20%. We train `Bio-JOIE` on this train set along with human PPIs. For evaluation, positive test samples and selected negative samples, mentioned in Section 3.1 are used to perform binary prediction. We adopt F1-score as the evaluation metric.

#### Results

As in Section 3.3, we first evaluate `Bio-JOIE` on SARS-CoV-2 PPI prediction. From the observation in Section 3.3, two important factors are considered: three aspects in the gene ontology domain and the scope of input SARS-CoV-2-Human PPIs. More specifically, we define increasingly four scopes of input PPIs, as shown in Figure 5, i.e. (1) S1: Only using the train folds of SARS-CoV-2-Human PPIs; (2) S2: Using SARS-CoV-2-Human PPIs with the 2-hop neighbor proteins from SARS-CoV-2 viral proteins, i.e. including the ones which also interact with any proteins that the SARS-CoV-2 interacts; (3) S3: SARS-CoV-2-Human PPIs with all other protein interactions on human; (4) S4: SARS-CoV-2-Human PPIs with all protein interactions in S3 plus all SARS-CoV and MERS PPIs. As for the aspects of the gene ontology domain, similar to Table 4 in Section 3.3, we adopt eight options, i.e. one without gene ontology information (NonGO), three using a single aspect of GO terms (BF, CC, MF), three options using two of the aspects (BF+CC, etc) and one using all three aspects (AllGO).

**Fig 5.**
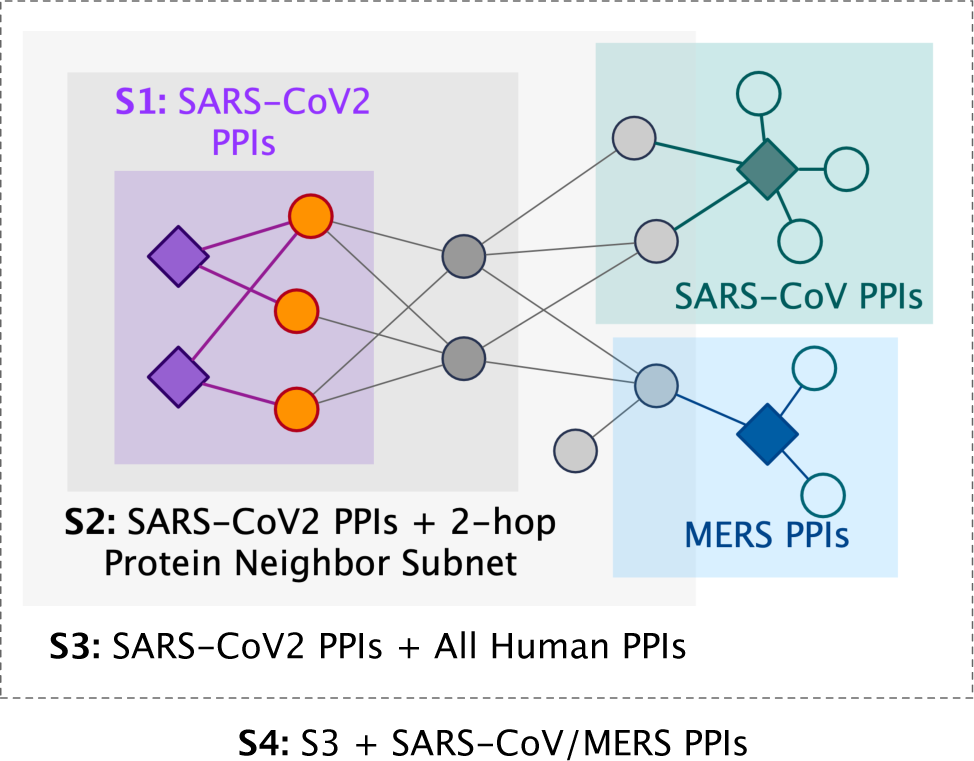
Different scopes of input to train `Bio-JOIE` for SARS-CoV-2 PPI prediction.

The results are summarized in Table 7. In terms of gene ontology aspects, we observe that CC contributes the most compared to other aspects of gene ontology annotations, and the best performance is achieved by adopting CC+MF in `Bio-JOIE` learning. One explanation is that most of the SARS-CoV-2 proteins have CC annotations and these annotations make up over 70% of all currently available annotations on average. However, less than 5 proteins (such as NSP and ORF 1a) have BF and MF annotations, possibly due to insufficient knowledge on understanding SARS-CoV-2 biological mechanism. As for the input fields, we find that the performance drastically increases with the expansion of input from S1 to S2, which indicates that interactions of 2-hop neighbor proteins can benefit SARS-CoV-2 PPI prediction. However, such a trend is not clearly observed when expanding the input field from S2 to S3. We hypothesize that proteins that are not within 2-hop neighbors may not be very related to SARS-CoV-2 or provide beneficial insights. Interestingly, when adding interactions of two related coronaviruses (SARS-CoV/MERS-CoV) that cause respiratory infection, the performance continues to improve with a relative gain of 3.4%. As shown in Figure. 2, viruses that are closely related to SARS-CoV-2 tend to share important properties. This strongly suggests that it is crucial to leverage their interactions and gene ontology annotations as augmented knowledge for drastically emerging SARS-CoV-2.

**Table 7.**
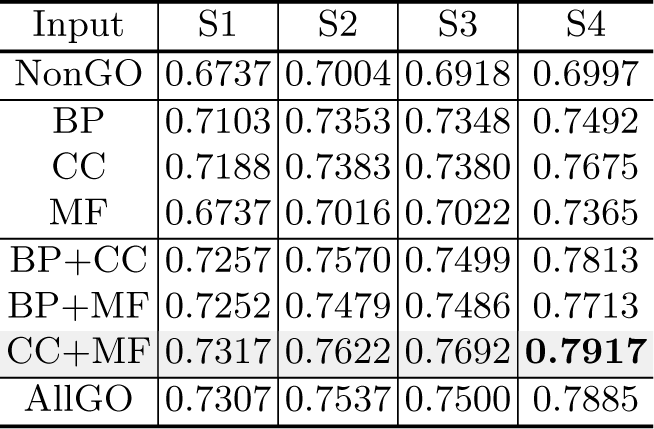
F-1 score on SARS-CoV-2-Human PPI interaction classification.

| Input | S1 | S2 | S3 | S4 |
| --- | --- | --- | --- | --- |
| NonGO | 0.6737 | 0.7004 | 0.6918 | 0.6997 |
| BP | 0.7103 | 0.7353 | 0.7348 | 0.7492 |
| CC | 0.7188 | 0.7383 | 0.7380 | 0.7675 |
| MF | 0.6737 | 0.7016 | 0.7022 | 0.7365 |
| BP+CC | 0.7257 | 0.7570 | 0.7499 | 0.7813 |
| BP+MF | 0.7252 | 0.7479 | 0.7486 | 0.7713 |
| CC+MF | 0.7317 | 0.7622 | 0.7692 | **0.7917** |
| AllGO | 0.7307 | 0.7537 | 0.7500 | 0.7885 |

Besides providing PPI prediction, the proposed model can help by identifying high-confidence candidates for potential human protein targets; this is considered as a link prediction task. When a viral protein (such as SARS-CoV-2 M protein) is given as the query, along with a specific relation (such as “binding” under the experiment system type of “Affinity Capture-MS”), `Bio-JOIE` can output a list of most likely protein targets by enumerating the triples with top *fr* (*h, t*) scores. The predictions are listed in Table 8. It is our observation, `Bio-JOIE` can successfully predict the high-confidence human protein targets in the test set from by [12] among its top predictions (marked as boldfaced entities). Other than the proteins in the test set, `Bio-JOIE` can also provide a list of reasonable candidates that possess a relatively high MIST score. For example, P62834 is one of the top-ranked protein targets of SARS-CoV-2 NSP7 by our `Bio-JOIE`, which has a MIST score of 0.658. Diving deep into the facts for P62834, though P62834 is not considered as a high-confidence target by [12], we observe that both P62834 (RAB1A HUMAN) and SARS-CoV-2 NSP7 interacts with protein P62820 (RAB1A HUMAN). Besides, they are both annotated with the cellular component GO:0016020 (membrane) and enables molecular function GO:0000166 (nucleotide binding), which are possibly the reasons for `Bio-JOIE` making such prediction with a high rank. Furthermore, `Bio-JOIE`’s predictions include proteins that are not covered by [12], which inspires further scientific research to verify.

**Table 8.**
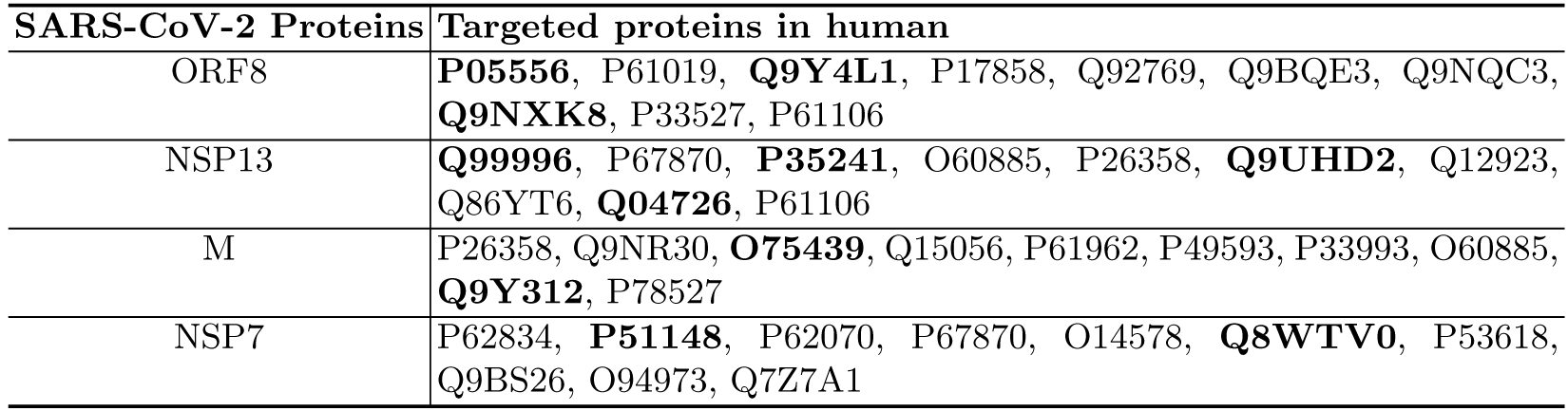
Top target proteins predicted by `Bio-JOIE`. Known interactions from training set are excluded. Proteins that are considered as high-confidence targets are boldfaced.

We further investigate how the information sufficiency of SARS-CoV-2 related PPIs in training set affect the performance. We define the train-set ratio parameter as means the proportions of the SARS-CoV-2-Human PPIs that are used for training `Bio-JOIE` and follow the aforementioned evaluation protocol on “NonGO/S3”, “CC/S3”, “CC+MF/S3” and “CC+MF/S4” as input other than the control of SARS-CoV-2-Human PPIs part. We plot the PPI results in Figure 6. As expected, when the proportion of SARS-CoV-2-Human PPIs used for training increases from 20% to 80%, the F1 score improves from 0.2-0.3 to around 0.8, which strongly confirms that the known SARS-CoV-2-Human PPIs serve as one significant factor to the PPI prediction. Moreover, the more knowledge we know about existing SARS-CoV-2 interaction, the more powerful the model is to predict SARS-CoV-2. We also observe that the performance is not saturated when the training ratio is approaching 100%, which possibly results from the fact that as a novel coronavirus, the current known interactions are still very limited. This encourages the scientific communities to unearth more knowledge on SARS-CoV-2; moreover, `Bio-JOIE` has the potential of bringing about significant advances based on new discoveries.

**Fig 6.**
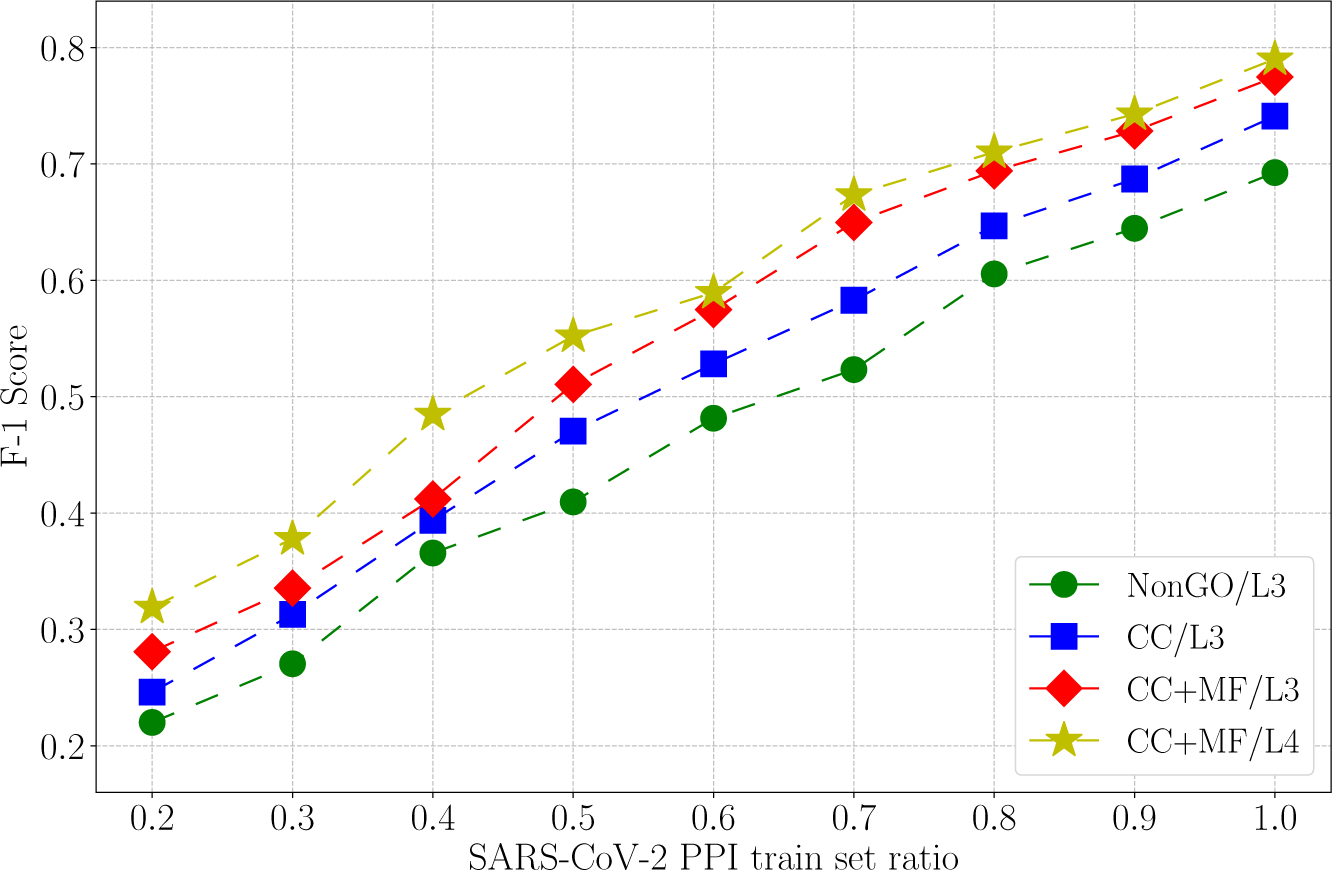
Bio-JOIE performance on different train-set ratios of SARS-CoV-2-Human PPIs.

## 4 Related Work

In the past decade, much attention has been paid to representation learning of KBs. Methods along this line of research typically encode entities into low dimensional embedding spaces, where the relational inference [32], proximity measures and alignment [6] of those entities can be supported in the form of vector algebras. Therefore, they provide efficient and versatile methods to incorporate the symbolic knowledge of KGs into statistical learning and inference. Some existing approaches focus specifically on computational biology studies [1, 9, 15, 27, 34], which similarly embed features of biological entities within low-dimensional representations. One representative work related to ours is Onto2Vec [27], in which protein representations are learned by incorporating the full semantic content of gene ontology in the feature learning using Word2Vec [22]. However, Onto2Vec replies on the ontology information, while falls short of capturing the multi-relational semantic facts that are important to characterize the proximity of biological entities. For example, regarding the protein and GO terms, the PPI knowledge and the non-hierarchical relationships between gene ontology entities (such as “regulates”) are not considered.

Another thread of related work is joint representation learning for multiple KGs, where embedding models are learned to bridge multiple relational structures for tasks such as entity alignment and type inference. MTransE [6] jointly learns a transformation across two separate translational embedding spaces based on one-to-one seed alignment of entities. Later extensions of this model family, such as KDCoE [7] and JAPE [28], require additional information of literal descriptions [7] and numerical attributes of entities [28, 31, 35] that are generally not available for biological KB. Our recent development on this line of research, i.e. JOIE [13] learns a many-to-one mapping between entity embeddings and ontological concept embeddings, and aims at resolving the entity type inference task using the latent space of the type ontology. One of the caveats is that JOIE does not specifically incorporate the specificity of concepts in the ontology in the transfer process, for which we find to be particularly beneficial in this problem setting. Besides, the aforementioned methods are mostly for general encyclopedia KBs (such as Wikidata, DBpedia) and have not been adapted for the purpose the modeling biological KBs. More specifically, in contrast to these methods, our method features the characterization of more complicated many-to-many associations between proteins and GO terms. Besides, instead of predicting the alignment of entities, we focus on transferring relational knowledge from one domain to enhance the prediction on the other.

## 5 Conclusion

In this paper, we present a novel model `Bio-JOIE`, that enables end-to-end representation learning for cross-domain biological knowledge bases. Our approach utilizes the knowledge model to capture structural and relational facts within each domain and motivates the knowledge transfer by alignments among domains. Extensive experiments on the tasks of PPI type prediction and clustering demonstrate that `Bio-JOIE` can successfully leverage complementary knowledge from one domain to another and therefore enable learning entity representation in multiple interrelated and transferable domains in biology. More importantly, `Bio-JOIE` also provides interaction type predictions on dramatically changing SARS-CoV-2 with human protein targets, which potentially brings reliable computational methods seeking new directions on drug design and disease mitigation.

In our main directions of future research, we plan to enhance and extend entity representations by systematically incorporating important multimodal features and annotations. For example, primary sequence information and secondary geometric folding features can be modeled simultaneously in protein networks and their combined representation can lead to a comprehensive understanding that will greatly benefit many downstream applications.

## Footnotes

1 Data source: https://coronavirus.jhu.edu/map.html, updated as 9pm (PT), 06/15/2020.

2 Data source: https://wiki.thebiogrid.org/doku.php/covid

3 Data source: http://geneontology.org/covid-19.html

